# The genetic equidistance and maximum genetic diversity hypothesis: Smoke and mirrors?

**DOI:** 10.1101/2023.02.14.528494

**Authors:** Yixi Zhang

## Abstract

As a novel molecular evolution model that was claimed to have an advantage over the molecular clock hypothesis^1–3^, the maximum genetic diversity (MGD) hypothesis was utilized to study the modern human origins^4^. Nevertheless, there are serious problems with this hypothesis and both it and its derivative studies should be treated with caution.

## Introduction

The advent and evolution of modern molecular phylogenetics had taken the evolutionary biology field by storm^5–8^. A widely accepted molecular evolution method is the molecular clock hypothesis proposed by Zuckerkandl and Pauling, directing phylogenetic analysis for over half a decade^9–10^. Recent establishment of a new molecular evolution theory, the MGD^1–3^, was claimed to substitute the molecular clock hypothesis by its presenters^3^, Shi Huang and his colleagues. The application of MGD to study modern human origins supports the East Asia origin^4^ rather than the Recent African origin which combines molecular and fossil evidence^11–13^. However, their criticism of the molecular clock mirrored their limited understanding of current molecular phylogenetics. Severe problems also existed in MGD hypothesis, and thus its applications are rather immature, risky and problematic.

Although there are many differences between these two theories, the most fundamental difference between them is that MGD suggests that there is an upper limit to genetic distance and genetic diversity, that multiple hits do not increase the genetic distance, and that macroevolution is the evolution of increasing species complexity, accompanied by a compression of the upper limit of genetic diversity in “complex” species^3^.

This paper will discuss the MGD’s refutation of the molecular clock and the reasonableness of the basic principles of MGD, while pointing out the problems that arise in its study in relation to modern human origins.

### Invalid refutations of the molecular clock

In 1962, Zuckerkandl and Pauling ventured that the number of differences between protein sequences could be used to infer the timing of species divergence, since hemoglobin amino acid sequence differences between closely related species were small and those between distantly related species were large. Mutations also seemed to accumulate at a constant rate^14^. In 1963, Margoliash compared hemoglobin sequence differences in yeasts, fish, birds and placental mammals and found it interesting that the number of amino acid differences between a closely related “complex” species and a distant “simpler” species is essentially the same^15^. This also holds true for several protein sequences^16^ and genomes^17^, a phenomenon that Huang refer to as “genetic equidistance”^1^. However, it is worth noting that their definition of genetic distance remains problematic, as will be discussed later. In 1965, Zuckerkandl and Pauling coined the term ‘molecular clock’ to refer to a theory of the kind they had previously described^18^, and Huang suggested that the molecular clock was inspired by the phenomenon of genetic equidistance in 1963^3^. However, there is a difference between the two explications, in that genetic equidistance emphasised the contrast between two closely related “complex” species and one distantly related “simple” species^15^, whereas Zuckerkandl’s 1962 paper emphasised that amino acid differences are greater the more distantly related the species^14^. Nevertheless, whether the molecular clock hypothesis was inspired by genetic equidistance or not, it didn’t disturb the discussion between the molecular clock and the MGD hypothesis, while the latter is actually inspired by genetic equidistance^3^.

The earliest molecular clock hypothesis suggested that the rate of mutation was constant and that the timing of divergence could thus be inferred from the number of mutations accumulated in a sequence^19^, and Kimura’s neutral theory that many loci in the genome are neutral and that variation at these loci does not affect the fitness of organisms^20^, thus exhibiting a more constant rate of mutation, which provides support for the molecular clock hypothesis. In 1973, Ohta proposed the nearly neutral theory, which argues that between neutral loci and selected loci, there exists loci that were weakly selected, so weakly that they are close to neutral^21^.

Kimura was also one of the pioneers of population genetics^22^, and with the help of the latter, the conflict between natural selection and neutral theory was gradually reconciled. It is now generally accepted that neutral loci conform to an evolutionary process called ‘genetic drift’^23–25^, which means that they are influenced by random factors. Evolution is the result of a combination of natural selection and genetic drift. The effective population size of a population thus becomes particularly vital, with small effective populations being more sensitive to genetic drift and large effective populations being dominated by natural selection.

One of the Huang’s refutations of the molecular clock (and the neutral theory) is the recent discovery of numerous functional regions among genomes^26–28^, which suggested the rare existence of neutral sites^3^. However, it should be stressed that the proportion of functional genome sites is still under fierce debate^29–30^ as well as the distinction between function and neutrality exists. Consider the question of whether the discovery that numerous non-coding regions of the genome are functional absolutely contradicts the neutral theory itself? Obviously not, because functional regions and neutral mutations are not incompatible, and they are selectionally neutral as long as they do not affect the fitness of an individual, since changes in molecular structure or even in the structure of organisms may not affect the fitness associated traits of organisms. A classic instance is the mutation of pink flower colour to produce a pale pink colour, which is still neutral once it does not affect a range of life activities including pollination, although it does affect the phenotype^31^. Thus functional regions can undergo neutral mutations and absolute non-functional genomic regions are not essential for the neutral theory to be valid, and previous study proposed numerous neutral mutations may be fixed in adaptive evolution^32^. In other words, gain of function mutations can also convert a neutral site into a functional site, even influencing the fitness of organisms. It should be stressed, however, that finding fewer neutral loci among genomes is beneficial for phylogenetic analysis, as neutral sites are more likely to be subjected to multiple hits, resulting in wrong tree topology^33^.

According to Huang’s works, phenomenon that many sites undergo multiple hits could deny molecular hypothesis, proving the introduction of the latter in modern phylogenetic analysis is misleading the entire evolutionary biology field^3^. Nevertheless, the problem of multiple hits caused by the limited character states (A, T, C, G, 4 states) of DNA in phylogenetic analysis had already attracted enough attention^34–35^. By introducing bootstrapping^36^ as well as a series of statistical inference to test the tree while utilizing Maximum Likelihood or Bayesian method rather than parsimony analysis^37^, phylogeneticists coped with this problem with various strategies. Ironically, the rather limited number of character states of DNA loci poses problems, but on the other hand also makes statistical simulations possible. The introduction of Markov model and other statistical approaches to establish substitution models could make the molecular phylogenetics more accurate^38^. Therefore, saturated sites could also be excluded during phylogenetic signal evaluation with the help of statistical approaches^34^. For instance, one of the most commonly utilized strategies was given by Xuhua Xia based on information theory^39^. In a sequence where no mutation occurs and all bases at all sites are identical, the frequency of particular base in the sequence is 1, which is the case of the minimum entropy. In a saturated sequence, the frequency distribution of bases at sites can be modeled by polynomial distribution. By comparing the observed entropy as well as other statistical parameters with the expected saturated sequence, the saturation of that sequence can be assessed.

Another important refutation of the molecular clock hypothesis claimed by Huang is the fact that mutational rates vary among genomic regions and clades^3^. However, the means of settlement had also already existed. With the introduction of Bayesian method and Markov Chain Monte Carlo, algorithm of molecular clock was evolving fast^40–43^. The substitution model assumes substitution rates of different bases, frequency, heterogeneity, ratio of unchanged sites and so on^40^. The phenomenon that distinct rates among regions and clades, named “heterogeneity”, could be coped by the applications of various models^34^. Phylogeneticists concern distribution of substitution frequency among sites. Substitution rates of different bases at one site also differ so there exists various substitution models such as F81^44^, GTR^45^ and JC69^46^. As for difference among sites, phylogeneticists utilized random variable like Gamma distribution to simulate it^40^. To sum up, advocators of the MGD hypothesis failed to provide a persuasive reason to deny the molecular clock hypothesis.

## Materials and methods

### Protein datasets

To examine the genetic equidistance at protein level, the author took partial materials from Yuan, 2017, while analyzing cytochrome c, one of the first evidence of genetic equidstance.

Protein sequences included in this paper (cytochrome c, Pre-mRNA processing splicing factor 8, Acetyl-CoA carboxylase 1 and U5 small nuclear RNP 200 kDa helicase) were downloaded from UniProt (http://www.uniprot.org).

### Nucleotide datasets

All the sequences were downloaded from NCBI (https://www.ncbi.nlm.nih.gov/gene).

### Alignment and distance measure

All the sequences were aligned using Muscle and ClustalW with default settings in MEGA v. 11, and amino acid pairwise distance was measured with Jones-Taylor-Thornton model, calibrating distance disturbed by multiple substitutions. For both sequences, rates among sites were modeled as Gamma distributed, assuming the parameter as 1.10 and 0.9. To assess the degree of substitution saturation in the Prpf8 coding region, the author plotted saturation curves for Prpf8 nucleotide sequences from 24 species, measuring their p-distance, transition distance as well as transversion distance while simulating the rates among sites with Gamma distribution and uniform rates.

### Phylogenetic analysis

Protein and nucleotide sequences of Pre-mRNA processing splicing factor 8 (Prpf8) were utilized to reconstruct the phylogeny of vertebrates involved in this work using Maximum Likelihood method.

## Results

Under the assumption that multiple substitutions would contribute to the increase of genetic distance in modern molecular phylogenetics, the consequences derived from the data (S1-S5) support that genetic equidistance is dispensable among clades. Phenomenon similar to genetic equidistance is observed when analyzing cytochrome c and Acetyl-CoA carboxylase 1, with the exception of *Mus musculus* and *Rattus norvegicus*, which are “simpler” than *Euungulata* while exhibiting the same distance when analyzing cytochrome c regardless of considering multiple substitutions, contradicting to the basic assumption of genetic equidistance. Phenomenons contradicted to MGD theory like both *Bos taurus* and *Ovis aries* are almost equidistant to human were also observed while analyzing Acetyl-CoA carboxylase 1 protein sequence. To obtain a more meticulous illustration of genetic distance, Prpf8 protein sequences of 21 species and nucleic acid sequences of 24 species were studied, and more evidence against genetic equidistance has been found from the results (S3-4). For instance, *Castor canadensis* are equidistant to human and *Mus musculus*, and genetic distance among species also undergo fluctuations, making the genetic equidistant phenomenon used by proponents of MGD hypothesis to dismiss the molecular clock even less reliable.

Mapping of the saturation curves for the Prpf8 coding region shows that this gene has not yet reached saturation, regardless of whether site heterogeneity is taken into account (Fig. 1 and S5). Such results refute the MGD hypothesis that the vast majority of loci in the genome had been reached their upper limit^3^. The phylogenetic tree was constructed based on Prpf8 protein and nucleotide sequences (Fig. 2).

**Figure 1.**
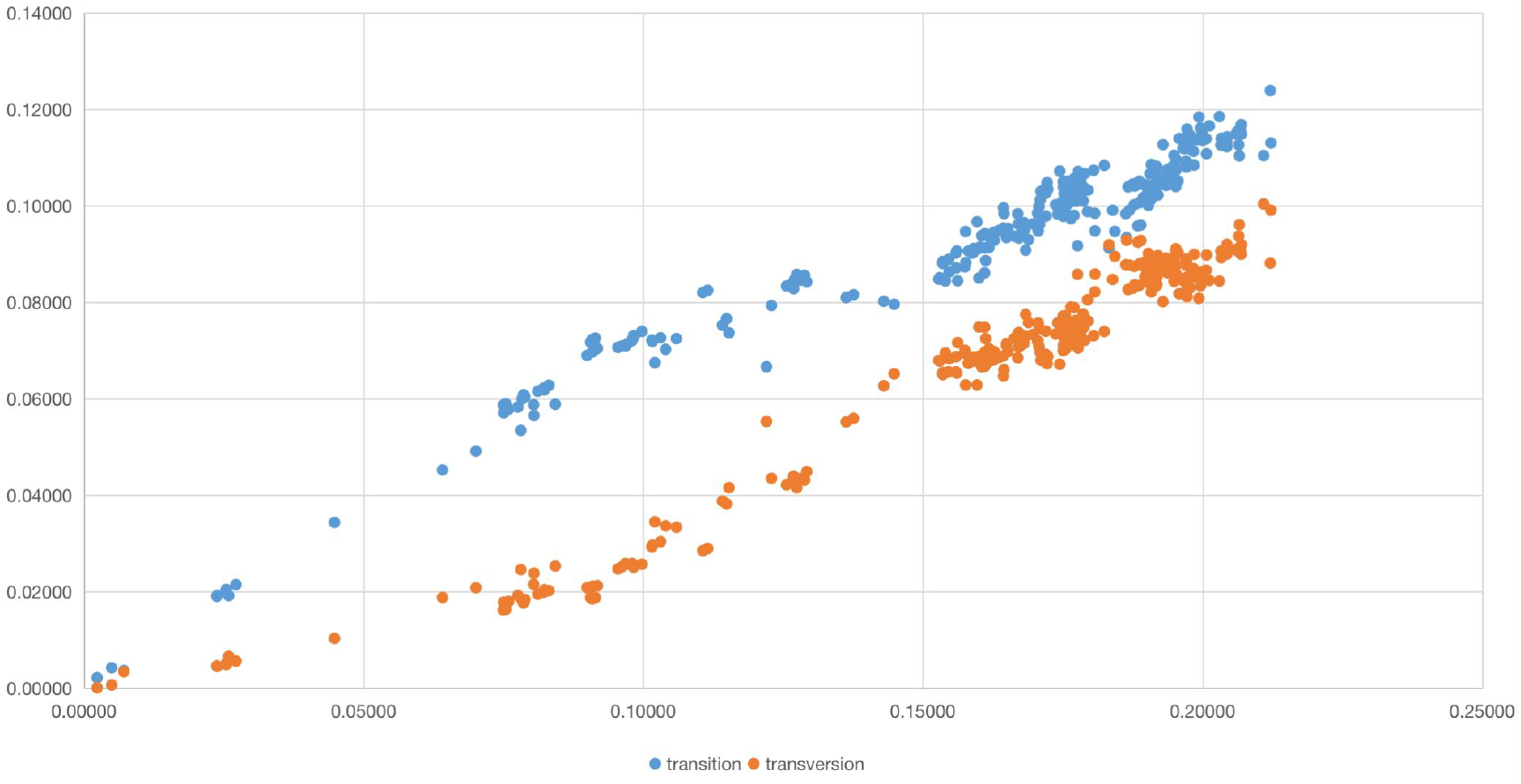
The saturation curves of Prpf8 coding regions. Results obtained under uniform rates are the same as that under Gamma distribution.

**Figure 2.**
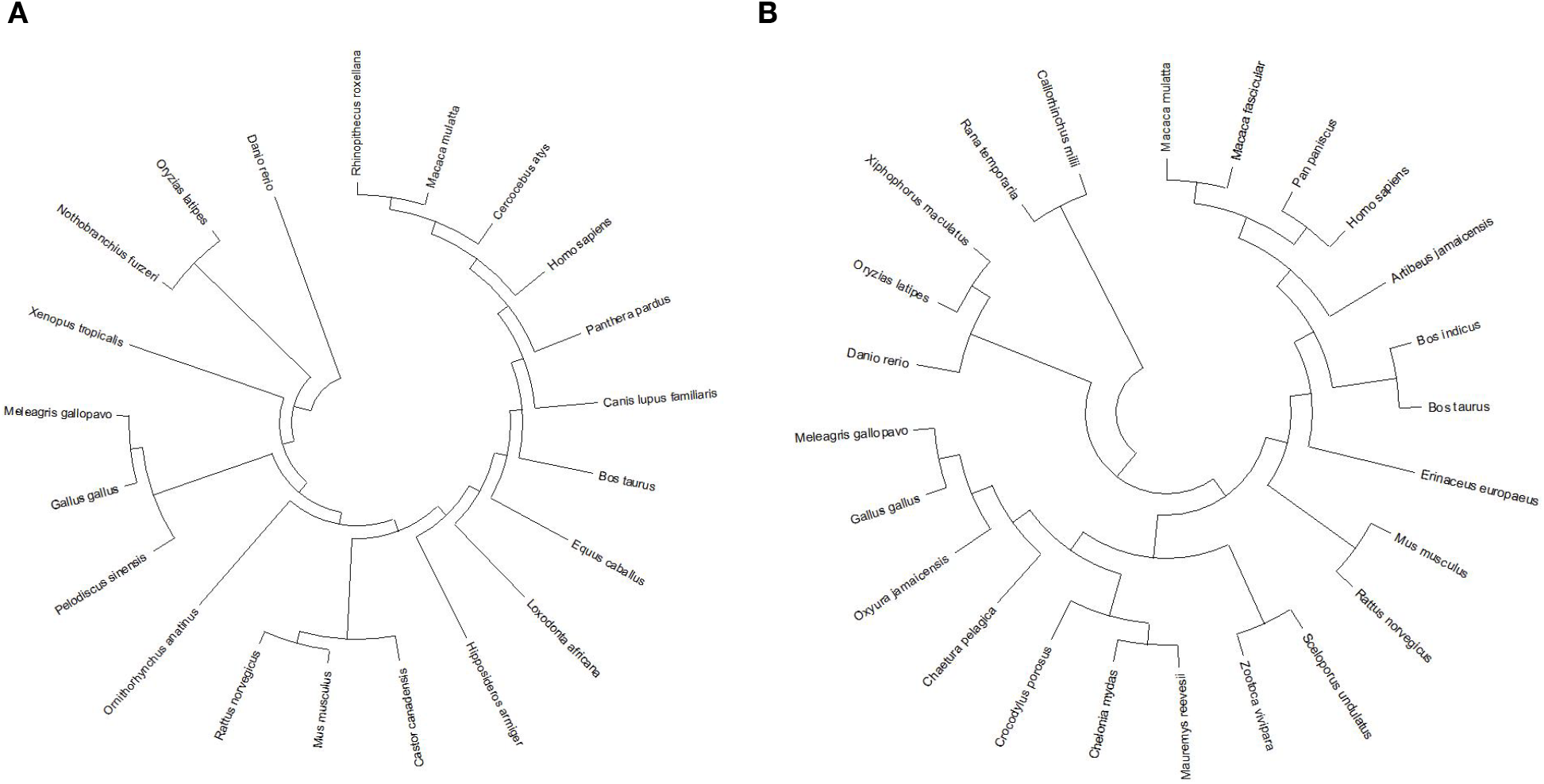
Maximum Likelihood tree constructed by Prpf8 protein (A) and nucleotide (B) sequences.

### What’s the upper limit theory?

Understanding the concept of upper limit of genetic diversity in the MGD hypothesis is rather simple. The MGD hypothesis suggests that multiple mutations at the same locus (multiple hits) do not cause any increase in genetic distance, then as more loci reach mutation saturation, genetic distance will converge to an upper limit^3^. Such an assumption would give rise to a serious problem called “long branch attraction” (LBA), which will be discussed later.

Of course the upper limit theory of evolutionary biology isn’t of no account, but it is a world away from the concept of upper limit of genetic diversity in MGD theory. Previous studies have demonstrated that essential genes which play a central role in metabolism, cellular processes, information processing and other life activities are highly conserved across species^47^. Chromosome-scale conservation of synteny among metazoan has also been reported, casting light on the evolutionary history of metazoan^48^. It has been well appreciated that the structure of several highly conserved proteins responsible for the maintenance of organisms will not change dramatically over time. Thus, in other words, to some extent, there exists an upper limit that the most basic proteins required for life will not change dramatically among organisms, so that some of the sequences in the genome are rather conserved. Nevertheless, the number of conserved elements reaching the upper limit should be fewer than all the conserved sequences observed. So if we look at genetic diversity at this large scale, then there exists a limit.

### MGD or LBA?

LBA is a frequent problem in phylogenetic analyses, leading to serious errors in the tree topology^34^, notably in molecular phylogenetic analyses. As mentioned earlier, the extremely limited number of character states is an inherent flaw of molecular systematics. Frequent substitutions in rapidly evolving clades will result in multiple hits that are misidentified as synapomorphies, allowing two unrelated long branches to be drawn together^49^.

Notably, the MGD hypothesis suggests that multiple hits and saturated loci will no longer lead to increased genetic distance^3^, so is the significant long-branch attraction the essence of MGD? The MGD method’s neglect of multiple hits is similar to the parsimony method’s branch length representing the minimum amount of evolutionary change, which is distinctively different from the Bayesian and maximum likelihood method’s branch length representing the expected value of each loci replacement^35^. Although the difference between MGD and LBA was discussed in Wang et al., 2020, its main content was to verify that slowly evolving proteins are closer to the neutral model and better suited for phylogenetic analysis. Only a short paragraph was focusing on the difference between LBA and MGD, presumably indicating that LBA is merely a special case of MGD. In Figure 4, they stated that 3-4 are saturated sites, and 5 is LBA^50^. In spite of the fact that MGD is indeed unequal to LBA, it still serves as a strong inducer of the latter, contributing to the distinctive consequences derived from the MGD hypothesis.

According to Huang’s comment^51^ on Ni’s article^52^, the more parsimonious (fewer ad hoc assumptions) a model is, the better it is. And in phylogenetic analysis, the Maximum Parsimony method is rather susceptible to disturbance by LBA^34^.

### Is macroevolution the evolution of species complexity increasing?

Despite minor differences in detail, the generally accepted definition of macroevolution is “evolution at the species level”^53^. MGD regards macroevolution as a process of increasing species complexity^3^. However, even regardless of the fact that the definition of complexity is inherently confusing, the evolution of complex traits in macroevolution is mosaic^54^. It is clear that at the scale of macroevolution, the complexity of distinct clades can reduce or increase by natural selection^55^. For instance, the untenable taxa Archezoa belonging to fungus that have lost their mitochondria, misleadingly simplified in structure, had been mistaken for “primitive eukaryotes before the acquisition of mitochondria”^56^. There are many other instances that complexity has been lost in macroevolution, especially the reduction in the number of cranial bones in amniotes compared to their sarcopterygii ancestor^57^ and the reduction in the degree of encephalization in some primate groups^58–59^.

